# Novel immune and stromal subtype classification system of lung adenocarcinoma based on tumor microenvironment

**DOI:** 10.1101/677567

**Authors:** Zihang Zeng, Jiali Li, Nannan Zhang, Xueping Jiang, Yanping Gao, Liexi Xu, Xingyu Liu, Jiarui Chen, Yuke Gao, Linzhi Han, Jiangbo Ren, Yan Gong, Conghua Xie

## Abstract

**Background:** Tumor microenvironment has complex effects on tumorigenesis and metastasis in lung adenocarcinoma (LUAD). However, there is still a lack of comprehensive understanding of the relationship between immune and non-immune stromal characteristics in tumor microenvironment.

**Patients and methods:** Eight cohort of 1681 lung caner patients were included in this study. The immune and non-immune stromal signatures of tumor microenvironment were identified by eigendecomposition and extraction algorithms. We developed a novel immune and stromal scoring system to quantify anti-tumor immune and promote-metastasis stromal activation, namely PMBT (prognostic model based on tumor microenvironment) as an R package. Tumors were classified into 4 subtypes according to PMBT system. Comprehensive analysis was performed in different subtypes.

**Results:** The 4 subgroups had different mutation landscape, molecular, cellular characteristics and prognosis, which validated by 7 data sets containing 1175 patients. 19% patients was characterized by highly active anti-tumor reaction with high production of immunoactive mediators, immunocyte, low fibroblasts infiltration, low TGF-β, VEGFA, collagen and glucose catabolic, low immune checkpoint per T cell, tumor mutation burden, and favourable overall survival (all, *P* < 0.05) named high-immune and low-stromal activation subgroup (HL). Cellular paracrine network showed both high humoral and cellular immune interaction in HL group. The low-immune and high-stromal activation group (19%) had opposite characteristics with HL group. Moreover, the PMBT system showed the value to predict overall survival and immunotherapy responses (all, *P* < 0.05).

**Conclusions:** Different molecular, cellular characteristics, mutation landscape and prognosis were discovered in the 4 subgroups. Our classification, PMBT system provided novel insight for clinical monitoring and treatment in LUAD.

## Introduction

Lung cancer is the most frequent tumor, of which non-small cell lung cancer (NSCLC) accounts for 80 percent [1]. Lung adenocarcinoma (LUAD) is well acknowledged for its malignancy and morbidity rate among people, and exhibits diversity of gene mutations. With the discovery of immune checkpoint inhibitors, such as programmed death 1 (PD-1) and its ligand PD-L1 (PD-1/PD-L1) [2], cytotoxic T-lymphocyte-associated protein 4 (CLTA-4), immunotherapy becomes a promising method to treat LUAD [3]. Nevertheless, immunotherapy only benefit ~16% LUAD patients for long-term survival [4]. It remains unsolved to deal with patients with therapeutic tolerance. Tumor stroma (non-immune) was reported to be closely related to the progression, metastasis and poor prognosis of tumors [5]. However, the immune and stroma components of tumor microenvironment (TME) are to be investigated. This study aimed to propose a novel insight and classification indexes to integrate both immune and stromal impacts on TME in LUAD.

TME is a very complex mixture, containing tumor cells, endothelial cells of blood and lymphatic vessels, fibroblasts, immune cells and normal tissues [6]. In the common analysis such as gene chip and second generation sequencing, the results may be unstable due to the influence of the TME. Therefore, this study tried to identify the characteristic genes of TME by signal decomposition algorithm. Non-negative matrix factorization (NMF) is a practical approach to decomposite matrix and distract abstract characteristics among massive data sets, which is commonly exploited for gene pattern recognition and computer vision in biomedical engineering [7]. Multitask learning (MTL) is an inductive transfer method to improve predictive capability by learning tasks in parallel while using shared representations [8].

## Materials and Methods

Eight data sets containing transcriptome expression profile and clinical information of 1183 LUAD and 498 non-small cell lung caner (NSCLC) patients (Supplementary Fig. S1, Supplementary Table S1) were included in this study. Unsupervised low-dimensional microdissected feature was extracted from TME by hierarchical clustering and non-negative matrix factorization (NMF) method [9]. Multitask learning was used to high Dimensionalization of features [10]. Consensus clustering and different gene expression (DGE) analysis were used to optimize gene signatures [11]. Immune and stromal scores were calculated by single-sample gene set enrichment analysis (ssGSEA) [12]. We proposed a scoring system that integrated immune and stromal scores to divide LUAD patients into the 4 subtypes, named PMBT which was encapsulated as a R package.

## Results

### The Identification of Novel TME subtypes in LUAD by PMBT System

In NMF analysis, we confirmed 4 TME-related NMF factors with different immune and stromal enrichment as reported previously (Supplementary Fig. 2B) [13, 14]. The enrichment analysis of the exemplar genes in high-immune and low-stromal activity NMF factor 6 showed high humoral immune response, T cell activation, response to IFN-γ (all, *P* < 0.000000001, *q-value* < 0.000000001, Supplementary Table S3, Supplementary Fig. 2D&E). In addition, activation of extracellular matrix organization, collagen fibril organization, response to *transforming growth factor beta* TGF-β (all, *P* < 0.0000000001, *q-value* < 0.0000000001, Supplementary Table S4, Supplementary Fig. 2F&G) were identified in low-immune and high-stromal activity expression pattern (NMF factor 5, Supplementary Fig. 2B).

The 179 TME-related genes were identified by MTL algorithm including many cluster of differentiation molecules and ligands, IFN-γ, *vascular endothelial growth factor A* (VEGFA), *integrin beta-2* (ITGB2), major histocompatibility complex molecule, chemoattractants and collagen related genes. Consensus clustering (K = 2) of the above 179 genes successfully divided the patients into the high immune-stromal ratio group and the low immune-stromal ratio group (Supplementary Fig. S4). Novel TME gene signatures were optimized using DGE analysis, and were divided them into 2 categories (108 immunity-related category genes and 58 stroma-related category genes, Supplementary Table S7, Fig. 1A). Novel immune and stromal scores were calculated by immunity-related and stroma-related genes by ssGSEA. The novel immune score revealed close links to immunocyte activation (Supplementary Table S8), favourable OS (*P* = 0.0038 in Cox regression) and low lymph node metastasis (*P* = 0.0315). However, stromal score was linked to cytoskeleton, collagen fibril organization, endothelial cell migration, glucose catabolic process to pyruvate, wound healing, VEGF, TGF-β (Supplementary Table S9), unfavourable OS (*P* < 0.0001) and high lymph node metastasis (*P* < 0.001), suggesting immune score reflected anti-tumor and stromal score reflected promot-tumor activation. Different immune and stromal activation divided the LUAD patients into the 4 subtypes (HL: high-immune and low-stromal; LH: low-immune and high-stromal; HH: double high; LL: double low). These 4 subtypes showed distinct TME-related gene expression patterns (Fig. 1B).

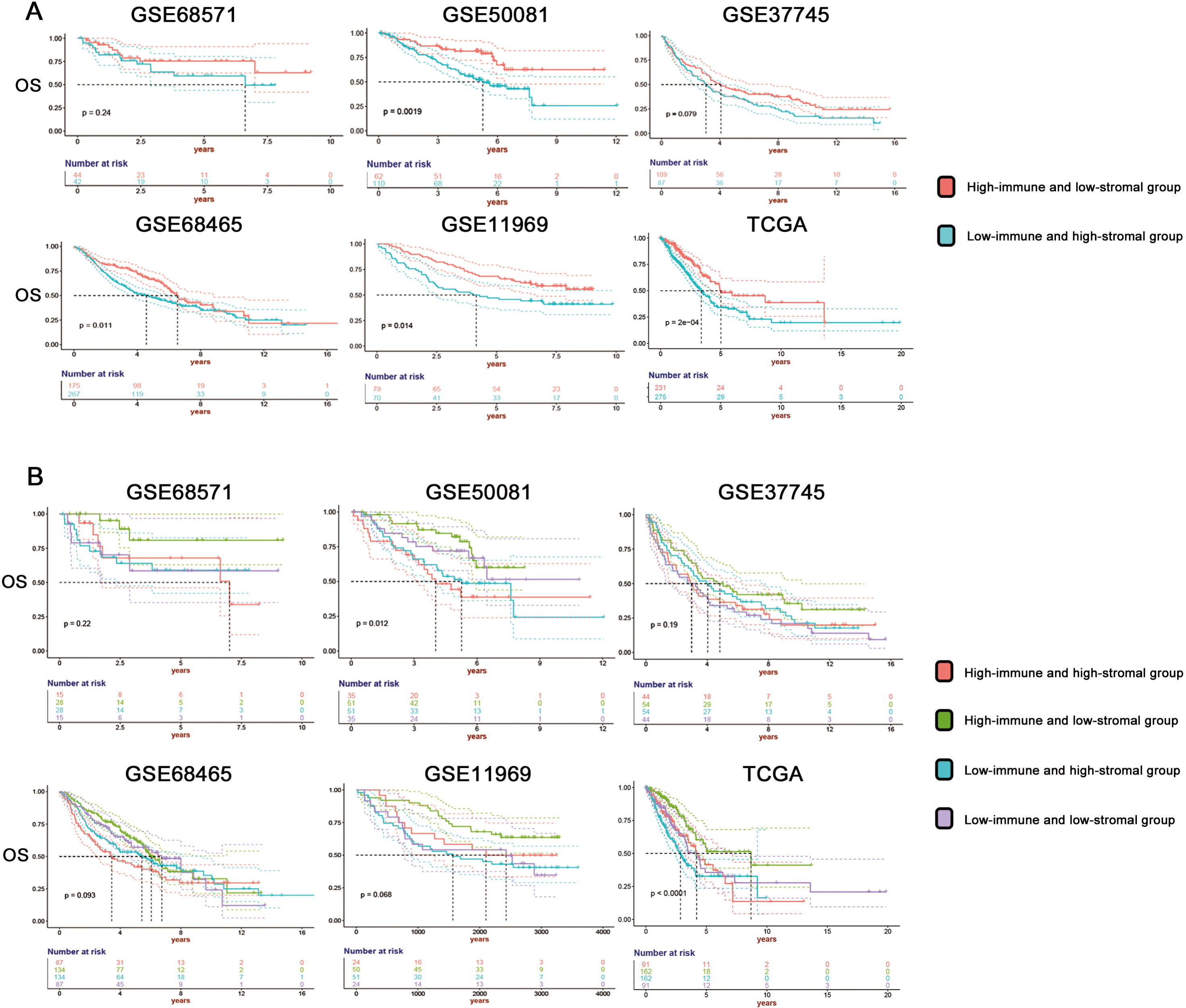

### Cellular and Molecular Characteristics in the 4 TME Groups Based on Immune and Stroma Related Gene Features

Extensive hyperimmune cell infiltration and TIL were found in the HH and HL class (*P* < 0.01, Fig. 1C). The LH and LL classes showed high fibroblast infiltration (*P* < 0.01, Fig. 1C). In the HH group, T cell activation, humoral immune response and extracellular structure organization (all, *P* < 0.000001, *q-value* < 0.01) were enriched (Fig. 1D, Supplementary Table S10). There were significant enrichment in the cytoskeleton, cell cycle regulation and DNA repair including microtubule cytoskeleton organization involved in mitosis and double-strand break repair in the LH group (all, *P* < 0.000001, *q-value* < 0.05, Fig. 1D, Supplementary Table S11). Significant immune activation containing B cell activation, T cell activation, interferon-gamma production and interleukin-12 production (all, *P* < 0.000001, *q-value* < 0.01, Fig. 1D, Supplementary Table S12) was observed in the HL group. No significant enrichment was found in the LL group (all, *q-value* > 0.05). Patients in HH and HL classes were related to more activation in immune molecules signaling, such as IFN-y, IL-1, IL-2, IL-6, IL-10 and PD-1 related signaling (*P* < 0.000001). However, HH had more PD-1 expression per T cell than HL (PD-1 expression/T cell abundance, *P* < 0.0001).

Six identified pan-cancer immune subtypes reported previously were integrated into our novel classification (Supplementary Table S13) [15]. We found that significant higher proportion of wound healing subtypes was shown in the LH group compared others (*P* < 0.000001), suggesting high proliferation rate, high expression of angiogenic genes and Th2 cell bias. HH group had high proportion of IFN-γ dominant subtypes (*P* < 0.000001) linked with high M1/M2 macrophage polarization, CD8 signal (Fig. 1E). Inflammatory class had a larger number than other groups in both HL and LL groups (*P* < 0.000001), suggesting low to moderate tumor cell proliferation, high Th17 and Th1 genes. The 2 low immune groups had more lymphocyte depleted subtype patients (*P* < 0.01) with Th1 suppression and high M2 response.

### Intercellular Communication Networks in TME

The core subnetwork containing CD8+ T cells, B cells, NK cells and fibroblasts was identified in all LUAD patients (Supplementary Fig. S8A). Different subnetworks were identified in different subgroups. Both stroma-centered subnetworks (fibroblasts, endothelial cells and laminin, Supplementary Fig. S8B) and immunity-centered (CD8+ T cells, B cells, CD molecule, interleukin receptors and TNF superfamily members, Supplementary Fig. S8C) were identified in the HH group. No stroma-centered and 2 immunity-centered subnetworks were identified in the HL group. One was cellular immune subnetwork consisting of CD4+ T cells, CD8+ T cells, NK cells, dendritic monocyte, C-C motif chemokine ligands and HLA-A (Supplementary Fig. S8D), and the other was humoral immune subnetwork containing B cells, neutrophils, macrophage monocyte, Class I major histocompatibility complex, leukocyte immunoglobulin-like receptors and C-C motif chemokine receptors (Supplementary Fig. S8E). Contrarily, in the LH group, only fibroblast-centered subnetwork containing ITGB4, LAMB1, LAMB3 and LAMC1 was identified in the LH group (Supplementary Fig. S8F). No subnetwork with more than 5 nodes was identified in the LL group, suggesting its desert-like molecular communication.

### Mutation Landscape and Tumor Mutation Burden of Different Immune and Stromal Class

Based on consistency clustering for the 166 TME characteristic genes, the low immune-stromal ratio group showed a higher number of mutations (*P* < 0.0001), higher tumor mutation burden (*P* < 0.0001) and a higher mutation rate and Variant allele frequency (VAF) in the driving genes (Supplementary Fig. S9A-D & 10A,). The low immune-stromal ratio patients had high TP53 mutations (55% vs. 38%, *P* < 0.001) with more frame shift deletion (13.46% vs. 8.60%, *P* = 0.341) and less nonsense mutation (16.03% vs. 22.58%, *P* = 0.2625). In addition to TP53, COL11A1 and KEAP1 also showed a large rate difference, and they had biological effects of extracellular matrix organization and MHC-mediated antigen processing and presentation in Reactome pathways database [16]. Moreover, higher mutation of WNT related genes was observed in the low immune-stromal ratio groups (Supplementary Fig. S10 B).

For the 4 subgroups, significant lower TMB were found in the HH and LH groups (*P* < 0.0001, Supplementary Fig. 9G). Moreover, high stromal score showed the positive correlation with TMB (cor = 0.2936, *P <* 0.00001, Supplementary Fig. 9H). Weak negative correlation between immune score and TMB was found in this study (cor = −0.1369, *P* < 0.01).

### The Clinical Characteristic of the 4 Subtypes was Significantly Different

The clinical characteristics of the 4 groups were shown in the Table 1. Pathological T stage (*β* = 0.4153, *P* = 0.0115) and tumor status (*β* = 1.0658, *P* = 1.48e-05) were retained as significant prognostic factors in multivariate Cox regression. Both of TNM stages and residual tumors were risk factors for OS (all, *P* < 0.01, Supplementary Table S14). Higher T and N stage was found in the LH and LL groups (*P* < 0.01), similar to N stage (*P* < 0.05).

The dichotomy of consistency clustering (the high an low immune-stromal ratio groups) showed significant prognostic differences of OS (HR = 0.5597458, *P* = 0.0004, Fig. 2A). For the 4 subgroups, HL patients had significantly better OS than the others (HR = 0.4617451, P < 0.0001, Fig. 2B). LH patients had worse OS (HR = 1.788947, *P* < 0.0001), and HH and LL patients were comparable (HR = 1.045836, *P* = 0.8443), despite their distinct TME. The median survival of the 4 groups had similar trend as OS (HL: 8.682192 years; HH: 4.194521 years; LL: 4.186301 years; LH: 2.857534 years).

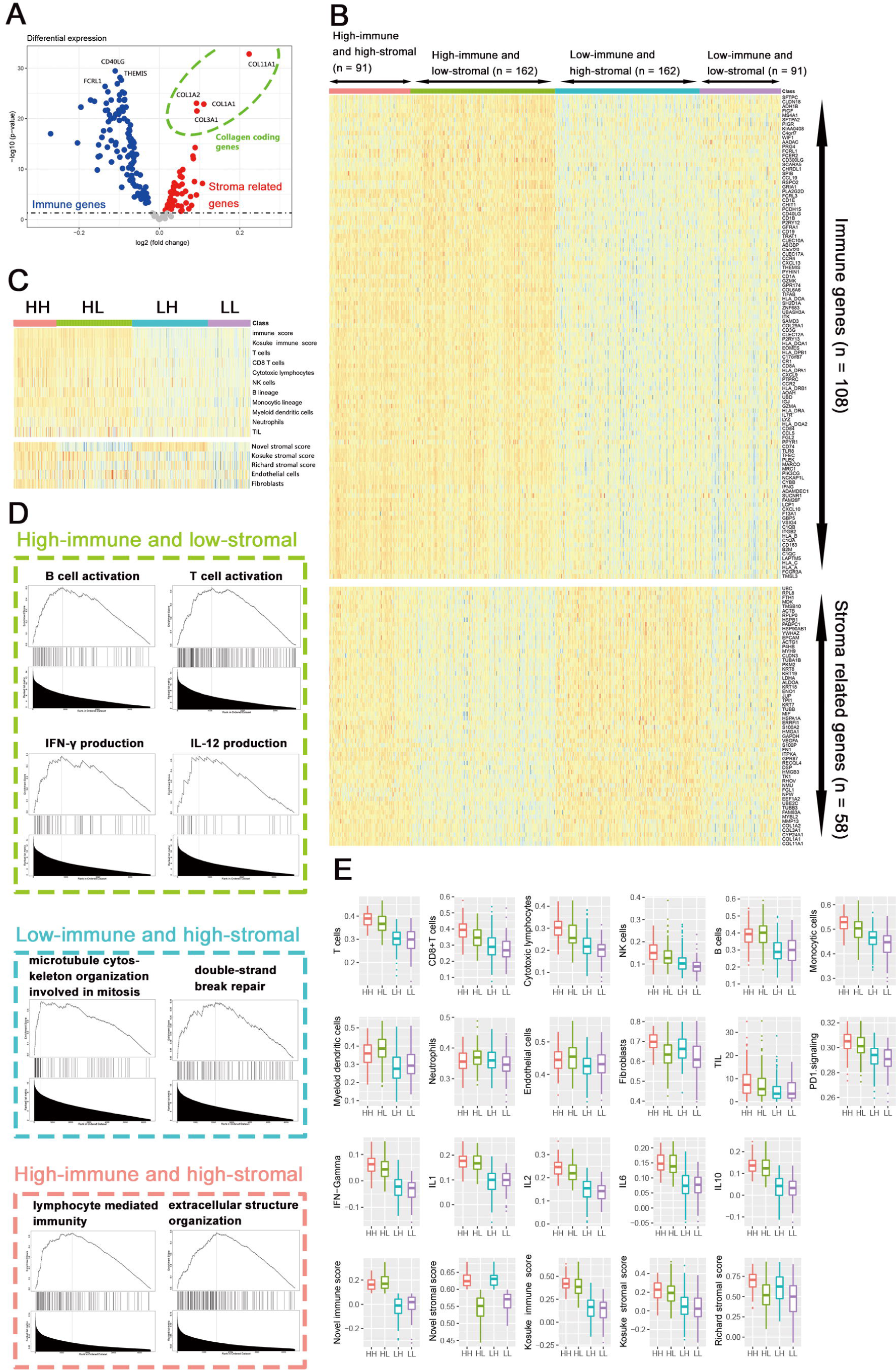

### PMBT Score System was Used to Predict Prognosis and Immunetherapy Response

PMBT scoring system consisted of immune score, stromal score and PMBT score, having closely relation with lymph node metastasis (all, *P* < 0.05), whch was validated by independent data GSE7339 (Supplementary Fig. 11A & B). Clearly, PMBT showed significant prognostic value for OS in all 6 data sets with survival information (Supplementary Fig. 12A, Supplementary Table S15). The effect of PMBT on prognosis were independent of the large rate difference and high mutation frequency genes in mutation analysis (TTN, KRAS, CSMD3, RYR2, LRP1B and ZFHX4, Supplementary Fig. 13). TP53 mutation linked unfavourable OS (*P* < 0.05), but hadn’t significant relation with the PMBT scores (Supplementary Fig. 14). Comparing with existing prognostic methods for LUAD, PMBT scores of 3,5-year survival were related to wider areas under the curve than other TME-based indexes (Supplementary Fig. 12B). In stroma data set (GSE22863), the 3 PMBT were significantly different in normal and tumor stroma (Supplementary Fig. 15).

Our novel PMBT scores was used to predict response in immunotherapy data set of melanoma [17]. While none of the 3 scores was correlated with anti-CTLA-4 treatment (*P* > 0.05, Supplementary Table 17), all the 3 scores of anti-PD-1 therapy showed a significant predictive effect in correlation analysis (all, *P* < 0.05, Supplementary Table 16). The subtypes significantly predicted the response of immunotherapy on anti-PD-1 therapy (HL vs. LH: 80% vs. 0%).

### Validation in Multidata Sets and Meta-Analysis

In all data with survival data (except GSE7339 and GSE22863), HH, HL, LH and LL had 19% (294/1551), 31% (481/1551), 31% (482/1551) and 19% (294/1551) patients, respectively. The validation data sets showed similar cellular and molecular Characteristics to training data (Supplementary Fig. S16), except GSE68571 (*P* > 0.05). The abundance of different cell population was comparable in GSE68571 (Supplementary Fig. S16), due to less mapping genes (5545 genes vs. more than 13,000 genes in other data sets).

In our bi-classification of NMF consensus clustering, 4 data sets with survival information showed significant better OS in the high immune-stromal ratio group (*P* < 0.05), and the 2 remaining data sets (GSE37745 and GSE68571) shared similar trend (Fig. 2A & B). Two-arm meta-analysis were performed for OS in all cohorts with prognosis information, and there was no obvious bias according to funnel plot asymmetry (*P* = 0.3014, Supplementary Fig. S17A). The high immune-stroma ratio group had significant better OS (HR = 0.63, 95%CI: 0.54-0.73, Supplementary Fig. S17B). Single-arm meta-analysis also showed that HL group had better 3 year (81%, 95% CI:73-88%, Supplementary Fig. S17C) and 5-year survival rate (66%, 95% CI: 56-76%, Supplementary Fig. S17D) than other 3 groups.

### The summary of the 4 TME subtypes in LUAD

The characteristics of the 4 subtypes are summarized in Table 2.

## Discussion

TME played an important role in tumorigenesis and development. Our study suggested more comprehensive understanding by considering both immune and stromal activity simultaneously. Virtual dissection of mixed tumor tissues was realized by signal decomposition algorithm NMF and other machine learning methods. We successfully identified the 166 TME related genes and constructed PMBT scoring system in LUAD. Based on PMBT, we classified LUAD into 4 subtypes with different molecular, cellular and prognostic characteristics.

Although the immune checkpoint inhibitor (anti-PD-1/PD-L1) treatment benefited NSCLC patients, only about ~16% patients had long-term survival under immunotherapy [18]. Screening of potentially sensitive population for immunotherapy helped to decrease medical expenses and improve quality of life. By calculating our PMBT system, we found that there was a significantly positive correlation among PMBT scores and immune response rates in anti-PD-1 therapy (all, *P* = 0.022). Moreover, HH had the highest expression of PD-1 and PD-L1 per T cell abundance, suggesting HH may benefit from immunotherapy (Table 2).

TMB have complex effects on tumorigenesis [19]. Driver mutations, such as tumor suppressor gene TP53, may increase genomic instability and increase cell proliferation, which links to unfavorable prognosis [20]. On the other hand, passenger mutations may activate the immune responses through the production of neoantigens and thus contribute to prognosis and immunotherapy response. Recent clinical studies revealed opposite prognostic effect of TMB in NSCLC patients without immunotherapy. In LACE-Bio-II (LB2) study [21], high TMB group (≥ 8 m/Mb) had better disease free survival (DFS), OS and lung cancer specific survival (LCSS) in 908 NSCLC patients, while the low TMB group (< 4 m/Mb) had worse prognosis (DFS: *P* = 0.007; OS: *P* = 0.016; LCSS: *P* = 0.001). However, another clinical study indicated that higher TMB (≥ 62 m/Mb) correlated with worse OS in 90 NSCLC patients (*P* = 0.0003) [22], especially in stage I NSCLC patients (OS: *P* = 0.0018; DFS: *P* = 0.0072). TMB may not be a very robust prognostic marker due to lack of elaborate consideration of the threshold, biological effects of individual mutation, driver and passenger mutations, as well as the interference of TME RNAs when sequencing. In our study, the HH and LH groups had significantly higher TMB (Supplementary Fig. 9G), suggesting TMB was related with both promote-metastasis stromal activation and anti-tumor immune activation. Single-cell whole exome sequencing may provide new insights due to higher purity of the tumor samples.

There are some limitations in this study: 1) lack of large sample size data for immune checkpoint therapy; 2) we did not set the uniform thresholds for 3 PMBT scores, due to heterogeneity of different cohorts; 3) low proportion of advanced stage patients were included in this study.

Overall, we identified both the immune- and stromal-related gene signatures in LUAD, constructed the novel scoring system and classified the tumor tissues into 4 TME subtypes with different molecular, cellular characteristics, mutation landscape and prognosis. PMBT showed excellent value to predict the prognosis and immune response. We expected more study to further validate our founding.

## Disclosure

The authors have declared no conflicts of interest.

## Acknowledgments

Thank Yuan Luo and Shijing Ma for their contributions to the writing assistance in this paper.

## Author contributions

CH-X and YG leaded and supervised the project. ZH-Z designed the analysis thought, and NN-Z and XP-J provided some points of view. JL-L and JR-C carried out data collection and standardization. YP-G, LX-X and XY-L collected various analysis tools. ZH-Z, JL-L, YK-G, LZ-H and JB-R contributed to statistical analysis. Codes and PMBT package were provided by ZH-Z. ZH-Z, JL-L, YG and CH-X wrote the manuscript.

